# Thunor: Visualization and Analysis of High-Throughput Dose-response Datasets

**DOI:** 10.1101/530329

**Authors:** Alexander L. R. Lubbock, Leonard A. Harris, Vito Quaranta, Darren R. Tyson, Carlos F. Lopez

## Abstract

High-throughput cell proliferation assays to quantify drug-response are becoming increasingly common and powerful with the emergence of improved automation and multi-time point analysis methods. However, pipelines for analysis of these datasets that provide reproducible, efficient, and interactive visualization and interpretation are sorely lacking. To address this need, we introduce Thunor, an open-source software platform to manage, analyze, and visualize large, dose-dependent cell proliferation datasets. Thunor supports both end-point and time-based proliferation assays as input. It provides a simple, user-friendly interface with interactive plots and publication-quality images of cell proliferation time courses, dose–response curves, and derived dose–response metrics, e.g. IC_50_, including across datasets or grouped by *tags*. Tags are categorical labels for cell lines and drugs, used for aggregation, visualization, and statistical analysis, e.g. cell line mutation or drug class/target pathway. A graphical plate map tool is included to facilitate plate annotation with cell lines, drugs, and concentrations upon data upload. Datasets can be shared with other users via point-and-click access control. We demonstrate the utility of Thunor to examine and gain insight from two large drug response datasets: a large, publicly available cell viability database and an in-house, high-throughput proliferation rate dataset. Thunor is available from www.thunor.net.

## INTRODUCTION

Understanding the effect of drugs and other perturbagens on cell proliferation has relevance to several fields in biomedicine, most notably in cancer (1)(2). Human cell lines provide a widely available, relatively standardized, and scalable *in vitro* system in which such effects can be quantified and compared (3). High throughput screening (HTS) is a framework in which cells can be imaged and counted at scale, across multiple cell lines, drugs and doses using large, robotically automated facilities, and more recently using all-in-one incubator and cell imaging devices e.g., IncuCyte S3 (Essen Bioscience Inc., Ann Arbor, Michigan, USA). In these studies, *in vitro* drug response is traditionally quantified in terms of cell *viability*, i.e., the cell count at a particular time point (usually 72 hours) post drug addition as a fraction of control (unperturbed) cells (4). Recently, we (5) and others (6) introduced novel drug-effect metrics based on cellular proliferation rates. These rate-based metrics avoid biases inherent in traditional viability assays, which can produce misleading interpretations, e.g., of cell line sensitivity to drug (5). While estimating proliferation rates is more demanding than performing traditional end-point assays, new automated systems (including the IncuCyte) have greatly reduced this burden. Therefore, for both end-point and time-based measurements, the primary challenge is no longer data generation but rather dataset management, analysis, and visualization at scale. Unfortunately, these tasks often involve a cumbersome and error-prone workflow involving processing of multiple instrument-exported file types, manual aggregation of spreadsheets, and analyses using (often costly) commercial software packages or custom code written in languages such as Matlab, R, or Python, which require time and computational skill to set up. Existing graphical software is often either specific to certain end-point only datasets (7) or lacks tools for annotating, storing and sharing datasets as well as interactive, multi-dataset visualization and statistics (8).

In this manuscript, we introduce Thunor (THOO-nor), a free software platform to address the challenges of analyzing and visualizing end-point and time-course cell proliferation datasets. We provide a description of the software and its web interface, inputs, and key features. We demonstrate the utility of Thunor with two case studies. First, we explore relationships between cell line drug sensitivity, drug pathway/molecular target, and tissue site of origin in the publicly-available GDSC dataset (9). Then, we demonstrate the use of proliferation rate-based data using an in-house, high-throughput proliferation rate screen, and show how Thunor can help check for common quality control issues and explore these data interactively. We then provide a brief discussion—methods and software implementation are described at the end. The Thunor software, documentation, and a read-only demonstration instance are available at www.thunor.net.

## RESULTS

### Software and web server description

Thunor is an open-source software platform that solves the storage, sharing, analysis, and visualization challenges of large-scale *in vitro* drug response datasets – both end-point viability and cell proliferation time courses. The drug-induced proliferation rate (DIP rate) (5) is a quantitative metric of cell proliferation calculated from time-course data; a set of values obtained from different drug concentrations can be fit by models of dose–response relationships and analyzed in an analogous manner to viability. To our knowledge, Thunor is the only tool that provides an interactive graphical interface for both types of data, combined with a database, group-based dataset sharing, and graphical annotation tools. A comparison to related software is provided in Supplementary Text S2 and Supplementary Table S1.

Thunor’s central features include a web interface drag- and-drop file upload with automatic dose–response metric calculation; a powerful, multi-paneled, interactive plot system with statistical analyses, inter-dataset comparisons, and a “tag” system which allows aggregation of drug and cell lines by categorical features of interest (e.g. drug molecular target, cell line phenotype, cell line tissue of origin etc.) for rapid, interactive, code-free analysis of large datasets (Fig. 1). Visualizations can be used for both quality control checks and analyses, and include automatic statistical tests where appropriate. Datasets and tags can be easily shared with other users by point-and-click.

**Figure 1.**
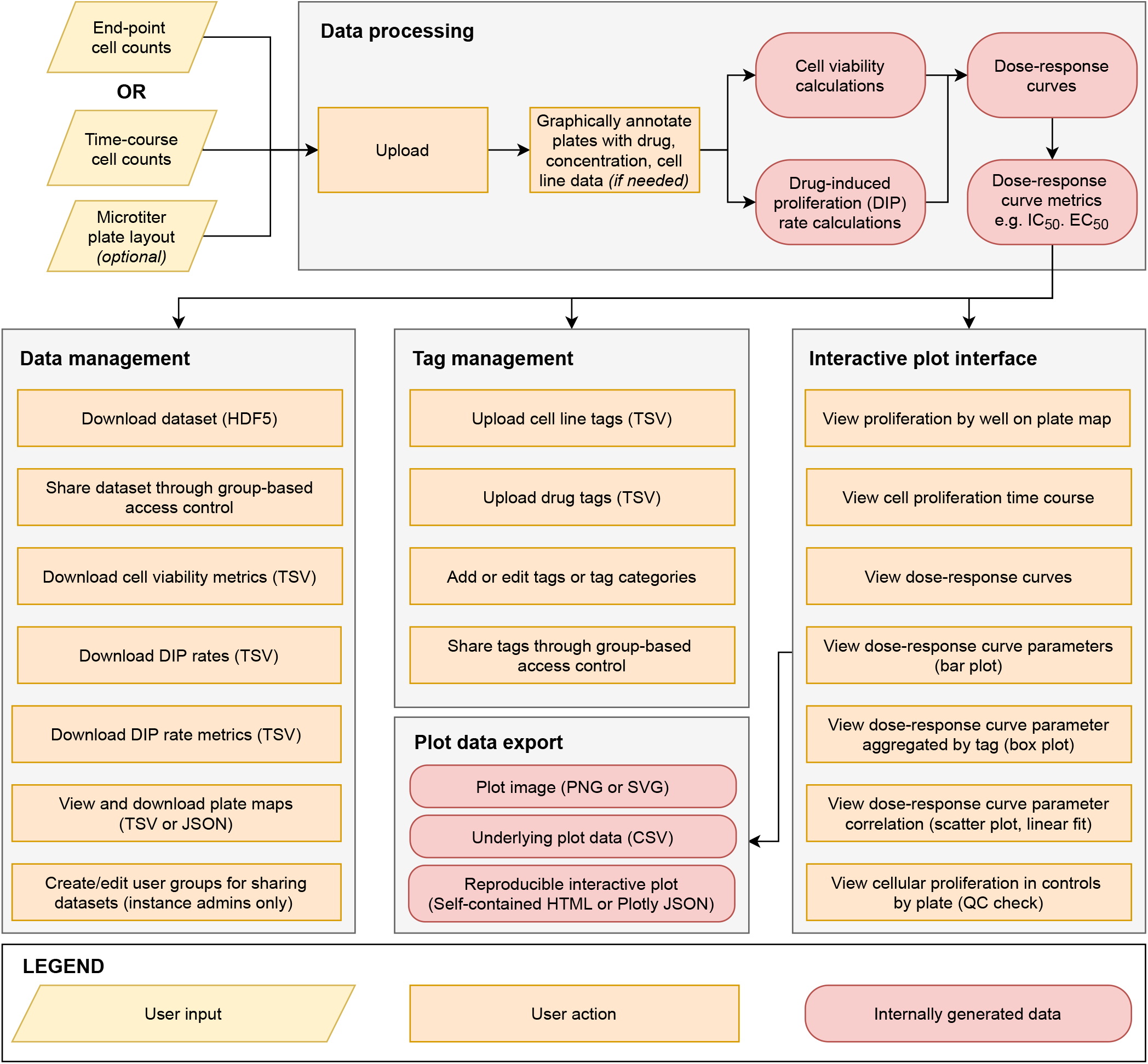
Thunor Web user workflow. Thunor accepts cell count data from end-point or time course experiments on microtiter (multi-well) plates. Layout data (describing cell line, drug, and drug concentration metadata) can be included or entered using a graphical interface. Data are automatically processed on upload and stored in a database, where they can be shared with other users, labeled with categorical tags for analysis, or explored using an interactive plot interface.

Thunor is split into a core Python library for analysis and visualization (Thunor Core) and a web application (Thunor Web). Thunor Core performs DIP rate calculation, curve fitting, and visualization capabilities. It can be used with Jupyter Notebooks (jupyter.org) and the wider Python ecosystem, enabling documented, reproducible analysis workflows to be archived and extended as needed. The Thunor Web interface accepts user data (cell count data; end-point or time-course), processes it into viability scores, DIP rates, and dose–response curves and their derived metrics using Thunor Core, and stores it in a database. These data can then be shared with other users, annotated with tags for aggregated analysis and statistics, and viewed and interrogated interactively (Fig. 2).

**Figure 2.**
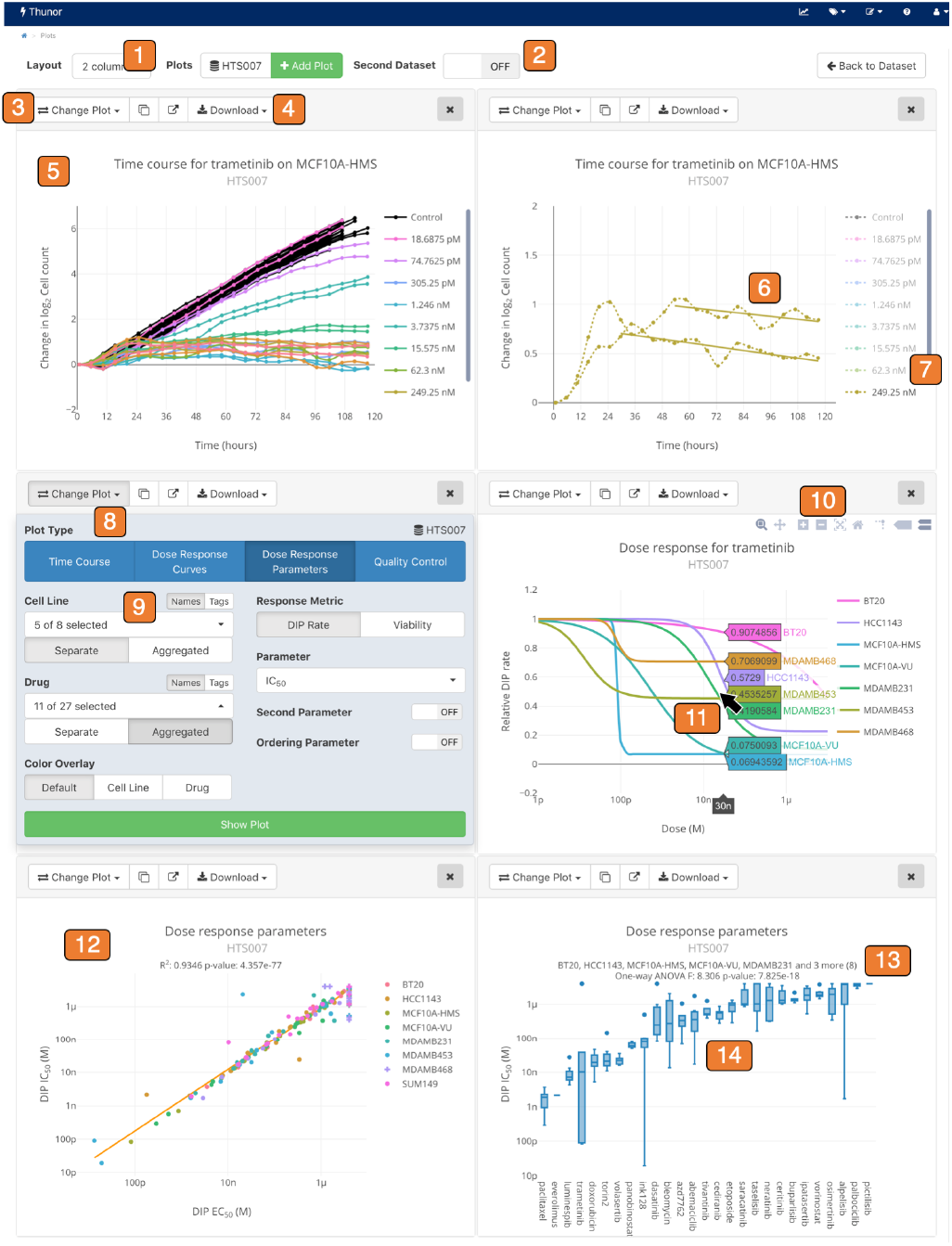
Thunor interactive plot interface. *1*. Multi-column layout option. *2*. Multi-dataset plot option. *3*. Plot toolbar. *4*. Download plot images (SVG, PNG) or data (CSV, JSON). *5*. Proliferation time course. *6*. Automatic DIP rate delay detection. *7*. Click to show/hide plot traces. *8*. Change plot panel. *9*. Flexible data selection and aggregation; by cell line, drug, or user-defined “tags.” *10*. Zoom, pan, and rescale axes. *11*. Hover mouse (tap on touch devices) to view underlying data. *12*. Comparison of two parameters (e.g., IC_50_ vs. EC_50_), or one parameter across two datasets. *13*. Built in statistical tests. *14*. Box plot showing aggregation of cell lines.

### Inputs

Thunor accepts cell count data in tab-separated value (TSV), HDF5, or IncuCyte Zoom (Essen Bioscience Inc., Ann Arbor, Michigan, USA) file formats from fluorescence-based or cell segmentation and counting platforms. Each contains a plate identifier, well identifier, cell count, and time point. The TSV format can be annotated (wells are already labeled with cell line name, drug name, and drug concentration) or unannotated. See Supplementary Text S1 for further details. For unannotated cell count data, the dataset can be annotated using a graphical plate map layout tool (Supplementary Fig. S2), or by uploading the annotation data in TSV or Javascript Object Notation (JSON) formats. The plate map layout tool can also export layouts in these formats for reuse on new datasets.

Tags are categorical data applied to cell lines or drugs, such as cell line tissue of origin, or drug class/molecular target. Tags can be uploaded as a TSV with columns “tag_name”, “tag_category”, and either “drug” or “cell_line” for drug/cell line tags respectively. Tags can also be entered and edited using Thunor Web’s graphical tag interface (Supplementary Fig. S3).

### Availability and documentation

Thunor Core and Thunor Web are freely available under the GNU General Public License version 3.0. An online, open access, read-only demo of Thunor Web is available at demo.thunor.net, which has been preloaded with the the case study datasets from this manuscript. A chat room provides an option to ask questions not addressed in the documentation and to contact the authors. These resources are all linked from the Thunor website, thunor.net.

### Case Study: Genomics of Drug Sensitivity in Cancer

The Genomics of Drug Sensitivity in Cancer (GDSC) (1) is a large dataset of cell viability and drug dose–response relationships. As an example of the utility of Thunor, we sought to identify drugs targeting cellular processes and pathways that have an outsized effect on collections of cell lines, grouped by their primary site/tissue of origin. Traditional analysis would either examine cell lines and drugs individually, or require custom code to group the data for analysis (9). Thunor enables these analyses from the graphical web interface.

Version 17a of the GDSC dataset (1) was downloaded and converted for use with Thunor HDF5 format (script included with Thunor Core). The converted dataset includes 72 hour luminescence-based cell viability assays for 1074 cell lines and 250 drugs over 9 concentrations. Annotation details, including cell lines’ primary site (tissue of origin) and drugs’ molecular targets and pathways, were similarly obtained, converted into a TSV file and loaded as Thunor tags.

Our GDSC analysis is summarized in Fig. 3. We examined a subset of cell lines from four primary sites (lung, breast, skin and aerodigestive tract; epithelial tissues known to be regulated by epidermal growth factor receptor (EGFR) family members (10)) for their sensitivity to drugs annotated by any of six cellular processes and pathways relevant to cancer (apoptosis regulation, cell cycle, EGFR signaling, ERK MAPK signaling, PI3K/MTOR signaling, and RTK signaling) using the activity area metric—the area above the dose response curve that increases with both drug efficacy and potency (Fig. 3A). As a group, cell lines from the aerodigestive tract had greater activity areas in response to inhibitors of EGFR signaling (Fig. 3B) and the drug afatinib appears to be driving the differences between cell lines from skin and the aerodigestive tract (Fig. 3C) and is further confirmed when examining the activity areas obtained from individual cell lines which identified TE-4, an aerodigestive tract cell line, as the cell line with the greatest response to afatinib (Fig. 3D). Afatinib and gefitinib, drugs that alter the activity of both EGFR and the related ERBB2, resulted in dose–response curves with relatively greater potency and efficacy compared to the other EGFR inhibitors (EGFRi) when applied to the TE-4 cell line (Fig. 3E). Investigation of the genetic alterations of TE-4 provided by the Broad Institute (11) uncovered genomic amplifications of both EGFR and ERBB2 (6 and 14 copies, respectively), suggesting a mechanism for its enhanced sensitivity. This analysis demonstrates how Thunor can be used to navigate a large dataset and focus down into a specific line of enquiry by following the data. Each plot can be produced in a matter of seconds, which enables rapid exploration of datasets with no programming needed.

**Figure 3.**
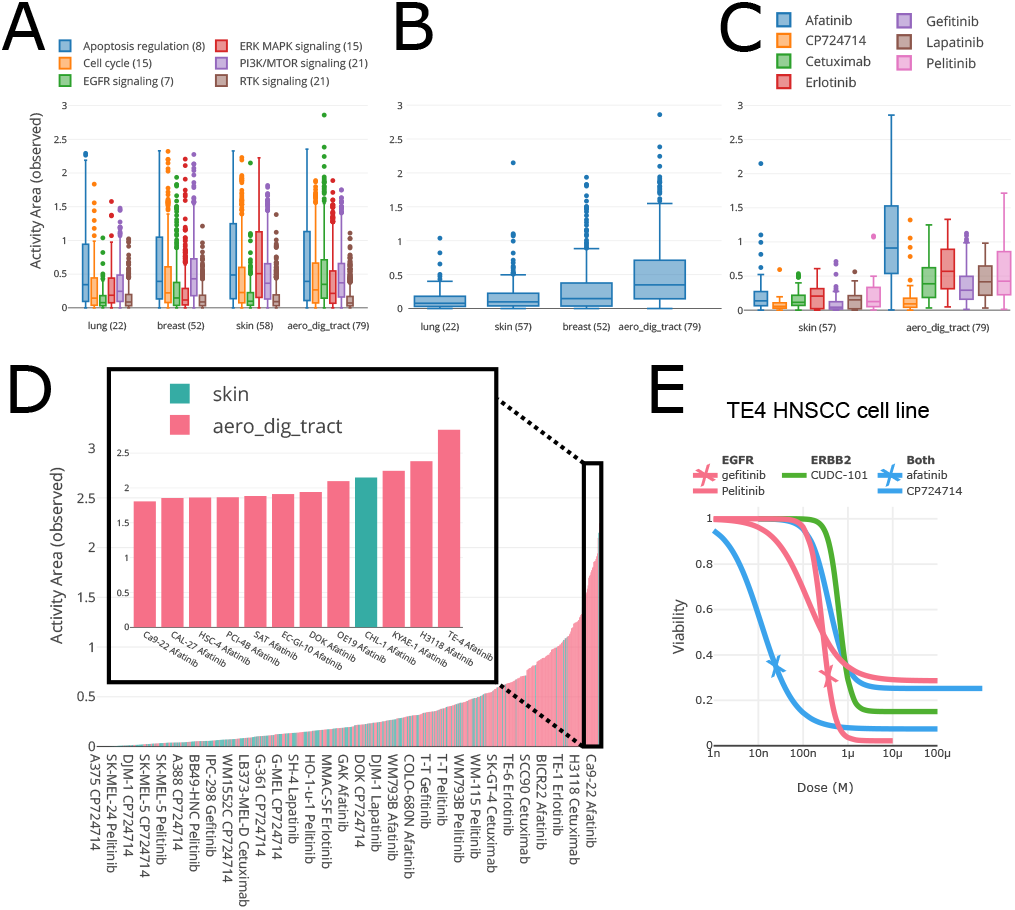
Thunor-generated plots for GDSC viability data. (**A**) Comparison of drug sensitivities between cell lines of tumors from different primary sites (lung, breast, aerodigestive tract (aerodig), and skin and treated with drugs of various classes, indicated by colors. The number of cell lines and drugs within each group are shown in parentheses and. Drug response is quantified by observed activity area (Activity area observed) and plotted as boxplots. (**B**) Only the data from (A) of EGFR-family-specific drugs are shown. (**C**) Only the skin and aerodigestive tract cell lines from (B) with responses to the individual drugs in the EGFR tag are shown. (**D**) Bar plot of sensitivity (AA observed) of cell lines from skin and aerodigestive tract cancers to EGFR-family-specific drugs. Colors correspond to cell line tags. Most sensitive cell lines have been expanded for easier visualization in the plot on *right*. (**E**) Dose–response curves of TE4 cells response to EGFR, ERBB2, and dual-targeting drugs respectively.

### Case Study: In-house cell proliferation dataset

The use of time-dependent measurements of cell proliferation instead of end-point viability can mitigate experimental biases found in the latter and has been shown to relate drug-response to cell phenotype (5, 12). Here, we demonstrate that Thunor can examine these data for common quality control issues and can visualize and analyze these data with the same ease as end-point data.

The HTS007 dataset contains a panel of eight breast cancer cell lines treated with 27 drugs at multiple concentrations generated in the High Throughput Screening Core of Vanderbilt University. Cell proliferation was quantified over five days. The dataset was uploaded to Thunor Web for analysis and visualization (Fig. 4). Thunor Web calculates the DIP rate and dose–response curves automatically, but it is useful to perform quality control (QC) checks to look for common experimental anomalies before proceeding. HTS007 used two plates for each cell line, thus we can check the reproducibility of cell growth rates in undrugged conditions across replicate plates for each cell line (Fig. 4A). We can also determine whether any spatial bias exists across the plate by visualizing the proliferation rate of each well overlaid onto the plate layout (Fig. 4B). This helps to identify, for example, if drugs were misapplied or the presence of “edge effects.”

**Figure 4.**
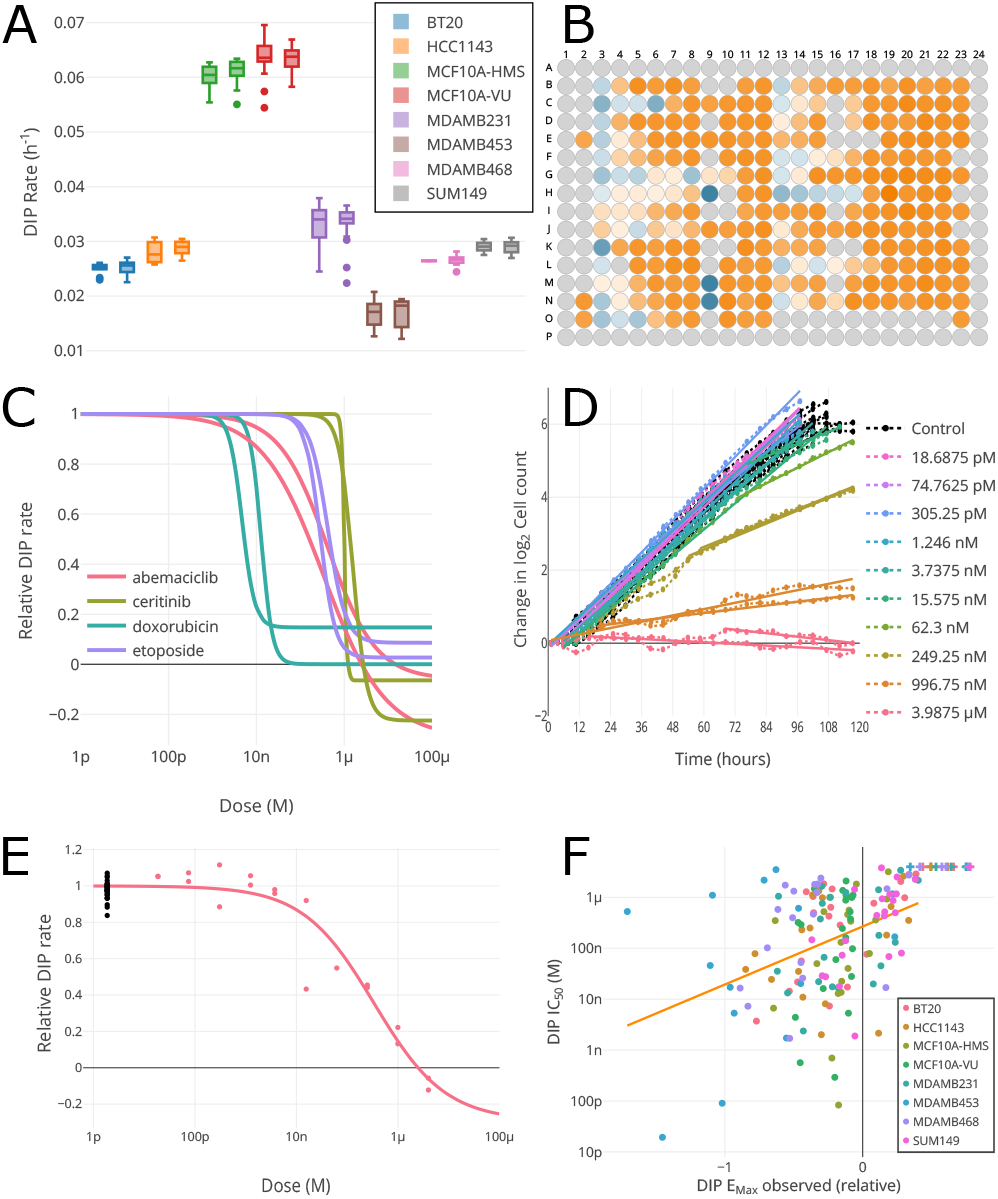
Thunor-generated plots for high-throughput time-course dataset HTS007. (**A**) DMSO control drug-induced proliferation (DIP) rates by plate, showing cell line growth reproducibility. (**B**) DIP rate by well (blue is negative growth, orange is positive, gray is no data). (**C**) MCF10A cell line relative DIP rates for four drugs. One of each color pair of lines shows data for MCF10A cultured at Vanderbilt (-VU); the other from cells cultured at Harvard Medical School (-HMS). (**D**) Cell proliferation time course (dotted lines) with DIP rate fit (solid lines) for each concentration replicated twice. The interactive interface allows trace hiding for added clarity. (**E**) Dose–response curve for MCF10A-VU cells in abemaciclib. Black dots show proliferation rate without drug (concentration for graphing purposes is set relative to lowest tested dose). Red dots show experimental replicates. Solid line shows model fit. (**F**) Potency (IC_50_) versus efficacy (maximum effect observed) for entire dataset, colored by cell line. + symbol indicates potency estimate truncated at edge of observed concentration range.

The HTS007 dataset contains two variants of the MCF10A cell line—one modified to express histone 2B conjugated to monomeric red fluorescent protein (H2B-mRFP) at Vanderbilt University (MCF10A-VU) and one expressing mCherry-conjugated H2B (H2B-mCherry) at Harvard Medical School (MCF10A-HMS). Data generation to evaluate the dose-dependent effects of drugs on cell proliferation rates was performed at Vanderbilt in both cases. The dose–response characteristics of the two cell line variants demonstrates high reproducibility, with similar dose–response curves obtained from four selected drugs (abemaciclib, ceritinib, doxorubicin, etoposide) (Fig. 4C). We confirmed that the dose–response curves are representative of the underlying cell growth characteristics at the various drug concentrations by visualizing the cell proliferation data (dash-dotted lines) and calculated DIP rate (solid lines, where the gradient is the DIP rate) for MCF10A-VU in abemaciclib at each concentration and well replicate (Fig. 4D). This view is useful for checking that any delay in drug response is correctly detected by the automated algorithm and the DIP rate quality of fit. Traces can be toggled on and off for clarity, where many concentrations are present. The DIP rate can also be viewed overlaid on the dose–response curve (fig. 4E), where each data point shows the DIP rate for each well replicate (corresponding to the gradients in fig. 4D). Black dots show undrugged/control data points, where the x-axis value is set one order or magnitude below the lowest measured dose due to the log scale. Thus, the quality of dose–response curves fits can be examined individually based on underlying data. Finally, as an example dataset-wide visualization, we show the maximum drug effect observed (efficacy) versus the IC_50_ (potency) for all cell line/drug combinations in the dataset, colored by cell line (fig. 4F). This shows how multiple metrics can be compared in Thunor; for example, one may wish to screen for drugs which are highly efficacious and potent. These plots are all interactive; one can hover the cursor (or tap on touch devices) to see information on data points and traces. Plots can be easily refined and altered in the interactive interface in a few seconds.

## DISCUSSION

High-throughput *in vitro* screens of large panels of chemical compounds against hundreds to thousands of cultured cell lines is a powerful and increasingly popular tool for probing complex intracellular networks and identifying druggable targets in hypothesis-driven biomedical research (13). There is a need for easy-to-use tools to rapid explore large cell proliferation datasets, including public datasets of end-point measurements and time-series cell count data, which are being generated in significant volumes by ourselves and other groups.

Although end-point data currently predominate over time-series data, a major Thunor feature is its ability to visualize and interpret both data types. We have been assessing the effects of drugs on cell proliferation over time for several years (5, 14, 15) and have promoted the use of the drug-induced proliferation (DIP) rate as a metric of drug effect (5). Another group has also promoted a rate-based metric of drug effect, the growth rate (GR; (6)), which utilizes a similar quantification approach. Although there is a GR web interface, GRcalculator (8), it lacks a database for storing and sharing datasets with granular permissions, does not have the variety of plot types available with Thunor, and lacks the ability to add and analyze additional annotation as with Thunor’s tagging system. Few other software packages have the ability to utilize time-series cell proliferation data. We provide an extended comparison of Thunor with this and other software in the supplementary information (Text S2, Table S1), but we believe Thunor adds significant value over the alternatives. Its interactive nature facilitates rapid exploration of any size dataset, and allows follow-up questions and hypotheses to be formed and investigated with a few clicks.

We anticipate that Thunor will stimulate collaboration between researchers, ease the exchange of drug-response data, and improve analysis reproducibility and transparency. Thunor is an active project and we encourage input and contributions from the research community. Extensions under consideration include drug combination response modeling, additional statistical analyses, improved quality control checks on data upload, and integration of -omics datasets (e.g., RNA-seq) to explore molecular correlates of drug sensitivity.

## METHODS AND IMPLEMENTATION

### HTS007 dataset

The HTS007 dataset (Data file S2) contains a panel of eight breast cancer cell lines treated with 27 drugs at multiple concentrations (four-fold dilutions). Each cell line was modified to express fluorescent histone 2B (H2BmRFP) to enable detection of nuclei via fluorescence microscopy. Cells were imaged by automated fluorescence microscopy approximately every four hours over five days in the Vanderbilt University High Throughput Screening Core. Nuclei were quantified by automatic image segmentation. The dataset is included in the online demo (demo.thunor.net).

### DIP rate calculation

The DIP rate is defined (5) as the gradient of the log_2_ cell count over time, after any initial stabilization period. The stabilization period is determined by iteratively excluding more time points from the beginning of the time course, evaluating goodness of fit at each step using linear regression, the root-mean-square error (RMSE) and adjusted R squared (ARSQ) are calculated. The final time point set is selected as follows:

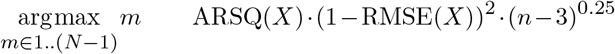

where *X* is the model fit, *N* is the number of time points available for the well, *m* is the index of first time point used for the fit, *n* = *N− m*+1 is the number of time points used for the DIP fit on the current iteration, and *fit* is a linear regression fit to the data points *m*..*N*. A minimum of two time points is required for a DIP rate fit (five or more is strongly recommended). When exactly two time points are present, the iterative procedure is skipped and the linear regression fit is used.

### Viability calculation

For viability calculations on multi-time point datasets, the closest time point to 72 hours is used. In the Thunor plot interface, the time point used can be verified by hovering the cursor over a viability data point in a dose–response curve. Viability is calculated as the ratio of the cell count in a well to the mean of the matched control wells’ cell counts at the same time point. Control wells are defined as wells on the same plate, using the same cell line, but with no drugs added to the well.

### Dose–response curves

Dose–response curves are fitted using a log-logistic function with three (viability) or four (DIP rate) parameters,

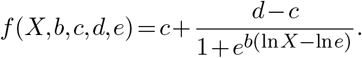

where *X* is a vector of concentration values and *b* (Hill slope), *c* (E_max_), *d* (E_0_), and *e* (EC_50_) are fit parameters. In the three-parameter case, *d* is set to 1 because viability is relative to control, i.e., the effect at zero drug concentration is, by definition, the control viability. The curve fitting is performed using the `curve_fit` function in SciPy (https://scipy.org/). Initial values for the fit parameters are estimated from the data using the same approach as the four-parameter log-logistic (`LL.4`) function in the `drc` R package (16).

In the DIP rate case, the `curve_fit` function selects a least squares fit using the Levenberg-Marquadt algorithm (17). The fit residuals are defined as

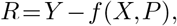

where *Y* is a vector of response values (i.e., DIP rate), *X* is a vector of drug concentrations, *P* is the set of fit parameters, and *f* is the log-logistic fit function defined previously. The standard error of DIP rate data points is incorporated into the fit by minimizing

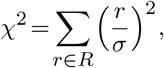

where *σ* is the standard error of a response value. Both control and experiment DIP rate values are used in the curve fit. Since the fit takes place in log_2_(concentration) space, a non-zero dose must be assigned to controls (log(0) is undefined). We set the concentration of controls to ten-fold less than the lowest concentration in *X*. The curve fit is replaced with a “no effect” model (shown as a horizontal dashed line in plots) if the dose– response curve is not significantly different from that no effect model (F-test, p*<*0.05). The fit is rejected (no dose–response curve shown) if any of the following occur: a numerical error occurs in the `curve_fit` function, the fit EC_50_ is less than the minimum concentration observed, or the fit E_0_ is greater than the mean plus one standard deviation of the control data points’ DIP rate values (where at least five control data points are used in the fit) or greater than 1.2*×* the mean of the control data points’ DIP rates (otherwise).

In the viability case, the sum-of-squared-residuals *R* is minimized directly. Dose–response curves are fitted with parameter constraints by the Trust Region Reflective algorithm (18). The parameter constraints are that Hill slope *b* must must positive, and E_max_ must be between 0 and 1 since viability cannot be negative.

The calculation of derived dose–response curve parameters like IC_50_ and activity area, and the available statistical analysis for different plot types, are covered in Supplementary Text S1. The difference between activity area based on the dose– response curve and activity area “observed” is shown in Supplementary Fig. S4.

### Thunor implementation

Thunor Core is a Python library, which provides core functionality, including structuring dose–response data and curve fit parameters using the `Pandas` library, automatically calculating DIP rate, fitting dose–response curve models, and plotting. Thunor Core can be used standalone, integrated into other processing pipelines, or utilized within Jupyter notebooks (https://jupyter.org), as shown in the Thunor Core online tutorial (part of the Thunor Core documentation, https://core.thunor.net). Thunor Web is built on Thunor Core, and is also written in Python using the Django web framework. It is deployed using Docker Compose, together with a PostgreSQL database, Redis database, and nginx web server. A script is included for easy deployment. An extended description of the software implementation and links to software dependencies are given in Supplementary Text S1; the architecture is shown in Supplementary Fig. S1.

### Software installation

Thunor Core is available from the Python Package Index (PyPI) with the command `pip install thunor` on Python ≥ 3.6.

Thunor Web is installed using Git (git-scm.com) and Docker Compose (docs.docker.com/compose). For convenience, a Python script is provided which automates the deployment process, including database initialization, creating an admin user, and adding transport layer security (TLS) encrypted connections, if desired, using Certbot (certbot.eff.org). Installation instructions are provided in Supplementary Text S3.

Both tools are compatible with Windows, Mac, and Linux. Smaller datasets (e.g., HTS007) require minimal resources; however for larger datasets like GDSC, a modern processor and 16GB RAM or more are recommended.

## ACKNOWLEDGEMENTS

We gratefully acknowledge technical assistance from J. A. Bauer at the Vanderbilt High Throughput Screening Core; M. Hafner and P. K. Sorger at Harvard Medical School for providing cell lines and drugs; C. M. Lovly for access to IncuCyte instrument and data; and C. E. Hayford, C. Meyer, and D. Westover for useful discussions. Funding was provided by the National Science Foundation (1411482 and 1942255 to C.F.L.), National Cancer Institute (U01CA215845 and U54CA217450, to V.Q. and C.F.L.), and Defense Advanced Research Projects Agency (Cooperative Agreement no. W911 NF-14-2-0022 to C.F.L.).

