## Supplementary Information for "Thunor: Visualization and Analysis of High-Throughput Dose-response Datasets"

Supplementary Material

#### Text S1: Extended description of Thunor implementation

##### Thunor Core/Thunor Web

Thunor software is split into Thunor Core and Thunor Web (Fig. S1). Thunor Core provides core functionality, including structuring dose–response data and curve fit parameters using the `Pandas` library, automatically calculating DIP rate, fitting dose–response curve models, and plotting. For data scientists, Thunor Core can be used as a standalone Python library on the command line, integrated into other processing pipelines, or utilized within Jupyter notebooks (jupyter.org), as shown in the Thunor Core online tutorial (part of the Thunor Core documentation, core.thunor.net). A list of key Python software libraries used by Thunor is included below.

| **Name** | **Role** | **URL** |
|---|---|---|
| `Bootstrap` | Front end layout | getbootstrap.com |
| `Certbot` | TLS certificate provisioning | certbot.eff.org |
| `Datatables` | Interactive web tables | datatables.net |
| `Django` | Web application framework | djangoproject.com |
| `Docker` | Application containers | docker.com |
| `Docker Compose` | Multi-container orchestration | docs.docker.com/compose |
| `Docker Machine` | Remote control and deployment | docs.docker.com/machine |
| `jQuery` | Front end interactivity | jquery.com |
| `Nginx` | Web server | nginx.org |
| `Numpy` | Numerical operations | numpy.org |
| `Pandas` | Data manipulation | pandas.pydata.org |
| `Plotly` | Graph drawing | plot.ly |
| `PostgreSQL` | Relational database | postgresql.org |
| `Redis` | Cache | redis.io |
| `Scipy` | Curve fitting, statistics | scipy.org |
| `Sentry` | Error aggregation, logging | getsentry.com |
| `uWSGI` | Application server | uwsgi-docs.readthedocs.io |
| `Webpack` | Static file bundling | webpack.js.org |

##### Thunor Core

Thunor Core is a Python package, which can be used interactively (Python prompt or within Jupyter Notebooks) or integrated into other data processing pipelines. Documentation is available at core.thunor.net. Thunor Core includes a suite of automated tests to help ensure code quality. Thunor Core is organized into several sub-packages: `thunor.io`, the core data types and file input/output; `thunor.dip`, which provides DIP rate calculations and derivative statistic calculations; `thunor.viability`, which calculates end point/single time point viability; `thunor.curve_fit`, for dose response curve fitting; and `thunor.plots`, for plotting; `thunor.helpers`, for miscellaneous "helper" functions. The core data structures are based around the Pandas DataFrame for high performance (McKinney, 2010). Numerical calculations and curve fitting use NumPy (numpy.org) and SciPy (van der Walt et al., 2011). Interactive plots are built using Plotly (plot.ly).

##### Thunor Web

Thunor Web is a web application built on Thunor Core and Django (djangoproject.com), a Python framework for web applications. Data are stored in a PostgreSQL database (postgresql.org) and accessed through the Django object-relational mapper. Redis (redis.io) is used as a cache to improve performance (optional). Thunor Web includes a suite of automated tests to help ensure code quality. Error monitoring, aggregation, and alerts are integrated with Sentry (getsentry.com). Sentry is available as open source software or a hosted service. Thunor Web incorporates user authentication. Account creation can be by open sign-up or by invitation. "Public" datasets can be available to view without authentication, or require a login. These preferences are set by the server administrator.

Thunor Web utilizes a "containerized" infrastructure based on the Docker framework (Merkel, 2014), which increases security and reproducibility. It also has a static file build tool, which bundles static files (those which don't change with every use, e.g. fonts, cascading stylesheets) into a cache-aware compressed bundle, which decreases bandwidth requirements and simplifies software updates. Thunor Web utilizes end-to-end transport layer security (TLS), which encrypts connections in the web browser for security and confidentiality.

#### Cell count file formats

Cell counts can be loaded into Thunor using one of three formats: Vanderbilt HTS Core format, Thunor HDF5 format, or IncuCyte Zoom format.

##### Vanderbilt HTS Core format

Thunor Web can accept annotated (with cell lines, drugs, and concentrations) or unannotated uploads in this format. Annotated files require all the fields in the table below, except those marked as optional. Unannotated files must omit all of cell.line, and "drug" prefixed fields - i.e. the only fields required for unannotated data are upid, well, cell.count and time.

If uploading unannotated data, plates can be annotated with cell lines, drugs and concentrations using Thunor Web's plate mapper.

Tab-delimited format, UTF-8 character encoding. Fields may be in any order. Extra columns may be present but will be ignored. The plate size will be detected based on the highest well number of the plate.

| **Field** | **Data type** | **Description** |
|---|---|---|
| `upid` | string | Unique plate ID (only needs to be unique within a dataset) |
| `well` | string | Well position on a plate (one character and two digits, e.g., `A01`) |
| `cell.count` | non-negative int | Cell count in specified well |
| `time` | non-negative float | Time (hrs) |
| `cell.line` | string | Cell line name |
| `drug1` | string | Drug 1 name |
| `drug1.conc` | non-negative float | Drug 1 conc. (molar) |
| `drug1.units` | string | Must be '`M`' |
| `drug2` | string | Drug 2 name (*optional*) |
| `drug2.conc` | non-negative float | Drug 2 conc. (molar; *optional*) |
| `drug2.units` | string | Must be '`M`' (*optional*) |
| `expt.id` | string | Experiment ID (*optional*) |
| `expt.date` | string | Experiment date (`yyyy-mm-dd` format; *optional*) |

##### Thunor HDF5 format

Files downloaded from Thunor, or from the Thunor Python package, in HDF5 format. HDF5 is a compressed, tabular, binary file format.

##### IncuCyte Zoom format

Thunor can also read files from the IncuCyte system from Essen BioScience. The IncuCyte Zoom software should be used to export a fluorescence marker proxy for cell counts. By default, the filename will be used as the plate name, unless a value is present in the Label: field.

The export can either contain one unified quantification per well (which by default is the median), in which case the header looks like the first example below, or each image can be exported separately, like the second example below. In the latter case, a unified score for each well is calculated as the sum of the values across all images at each time point.

Example header 1 (one count per well):

```
Date Time Elapsed A1 B1 C1 ...
```

Example header 2 (multiple counts per well):

```
Date Time Elapsed A1.Image1 A1.Image2 B1.Image1 B1.Image2 ...
```

#### Tag system

Thunor Web provides an interface for creating, editing, viewing, and deleting "tags," labels that are used to group sets of cell lines or drugs in the plot system. Tags can be created manually or uploaded from file and can be private, shared within a group, or public. Typical tags include cell line mutations, cell line tissues of origin, drug molecular targets, and drug classifications. Thunor's tagging feature is rare among similar open-source and commercial software tools (Table S1) and significantly aids in the visualization and analysis of large-scale datasets. An example set of tags from the Genomics of Drug Sensitivity in Cancer (GDSC) dataset (Garnett et al., 2012) is included in Data file S1.

#### Plate map designer

High-throughput experiments generally involve multiple plates with differing drugs, concentrations, and cell line layouts. Annotated plate maps are crucial for linking cell counts back to experimental conditions but are often missing from instrument-exported data. Thunor accepts cell count uploads (TSV format; see below) with or without plate map layouts. In the latter case, a user-friendly graphical interface (Fig. S2) is provided that allows users to manually annotate plates, as well as visualize pre-annotated datasets and export plate map annotations to file (TSV and JSON formats). Users can select an individual well on a plate and enter the cell line, drug(s), and concentration(s) used (drug combinations are supported). Cell line and drug name suggestions are provided from the database; the user may also create new ones. The plate mapper also has numerous features that drastically speed up data entry, including an "auto-stepper" that moves the current well selection (e.g., one well to the right) after data entry, saving keystrokes or cursor movements, an "auto-dilution" feature for changing concentrations using the auto-stepper, and a "template" system for annotating multiple plates simultaneously with the same details (e.g. for multiple plates with the same cell line).

#### Containerized infrastructure

Thunor Web provides a complete configuration set up using Docker (Merkel, 2014) and Docker Compose. After Docker is installed and Thunor Web has been configured, the user can build and start the application, web, and database servers with a single command: `docker compose up -d -build`. A "quick start" configuration is provided. Configuration is entirely managed by setting environment variables, as recommended by the influential software engineering guidelines "The Twelve-Factor App" (12factor.net/config).

The use of Docker containers enhances security (containers are isolated and firewalled), transparency (build specifications are in human- and machine-readable plain text), and reproducibility (specifications are standardized and work the same across different host systems). Docker Compose allows services to scale to incorporate larger installations behind a load balancer and is cross-platform. Thunor Web can be deployed to the local computer (on which the installation is run) or remote servers (including cloud providers) using Docker Machine (docs.docker.com/machine/overview). Installation of Thunor Web using Docker Compose and Docker Machine are described in the documentation (docs.thunor.net).

##### Static asset build system

Static assets are files that do not change depending on who requests them (e.g., JavaScript, cascading stylesheets (CSS), fonts). Complex web applications often require multiple such files for each page load, which can be slow. In addition, assets can get cached by the user's browser, making timely updates more difficult (the cache becomes "stale"). This can lead to errors if, for example, the user loads an updated version of a web page and an older JavaScript file is retrieved from the cache—the two files are not designed to work together. Thunor Web has a static file build tool, which uses Webpack (webpack.js.org) in a Docker container. Webpack combines multiple files into a single bundle, which is "minified" (the file size is reduced by removing whitespace and unused code) and compressed. Each bundle has a unique hash code attached to its filename, which avoids the stale cache issue described above (so-called cache busting).

##### Automated setup of TLS encryption

High throughput data are often proprietary or sensitive. Without encryption, all data transmitted between the web server and the user, including passwords, could potentially be monitored and intercepted. Even with open datasets, the protection of login details and user privacy remains important. These issues can be mitigated by use of transport layer security (TLS), which provides end-to-end encryption between the user's web browser and the server. However, TLS can be challenging to obtain and configure. Most certificate authorities do not issue certificates for free, and the alternative, self-signed certificates, are subject to forgery and generate warnings in the user's web browser.

Thunor Web provides a Docker-based wrapper around the Certbot tool from the Electronic Frontier Foundation certbot.eff.org. Certbot generates TLS encryption certificates for free. The Thunor Web Certbot wrapper generates certificates for the user's domain or subdomain where they host Thunor Web. The wrapper also uses OpenSSL to generate Diffie-Hellman parameters, which are used to create ephemeral keys for "forward secrecy." We also provide an example configuration file for the nginx server used by Thunor Web. The configuration has an A+ rating on Qualys SSL Labs' test (ssllabs.com/ssltest) at the time of writing.

##### Derived dose response parameters

In addition to the fit parameters defined in the main manuscript, Thunor also calculates numerous derived dose–response parameters. These include inhibitory concentrations (IC values), effective concentrations (EC values), and activity area (AA). The two variants used in this manuscript are the $IC_{50}$ and observed AA.

$IC_{50}$ is the half-maximal inhibitory concentration:

$$I_{0.5} = C_{0.5} \left( \frac{1}{1 - 2E_m/E_0} \right)^{1/h}. \tag{1}$$

Activity area is defined as the area above the dose response curve, bounded by the minimum and maximum experimentally tested concentrations. Thunor provides two variants: activity area (AA), which calculated activity area based on the fitted dose response curve, and activity area observed ($AA_{observed}$), which calculates the area between the dose-response curve (Supplementary Fig. S4).

##### Statistical calculations

Within Thunor's plot interface, several statistical tests are automatically performed (calculated using SciPy). On scatter plots, a line of best fit is calculated using linear regression and shown as an orange line, with the $R^2$ and p-value for the fit shown above the plot. On box plots, one-way ANOVA is used to test for a statistically significant difference in means of a dose response–parameter across a set of user-defined groups (using Thunor's tag system). Depending on the groups used for the box plots, an ANOVA test may not be statistically appropriate—it is left to the user to determine whether it is so. In particular, one-way ANOVA assumes that (1) response variable residuals are normally distributed, (2) population variances are equal, and (3) responses for each group are independent and identically distributed (i.i.d.) and normally distributed random variables. On bar plots, if the criteria described here are met, a two-sided Mann-Whitney U test is calculated, which tests for statistically significant difference in the ranks of two random variables. Here, Thunor will test for the difference in the ranks of a dose response parameter (e.g., $IC_{50}$) between two groups (cell lines, drugs, or tags). The test will only be run if the plot contains exactly two groups (defined by the number of cell lines/drugs/tags and the "color by" option, which defines the grouping variable), and both groups contain more than 20 data points. The test has continuity correction applied and assumes that data points are independent.

#### Text S2: Related software

A variety of open-source and proprietary software platforms/packages exist for analyzing and visualizing cell proliferation datasets. Proprietary packages generally contain more features and are more user-friendly, but require purchase of a product key or site license, which can be expensive.

Free and open-source tools are obviously cheaper and tend to be more flexible but require more technical expertise to use (e.g., programming experience). Moreover, most existing tools exclusively support end-point viability data, although more recent packages have been developed that include proliferation rates (i.e., time-course data). Thunor attempts to address the shortcomings of both free and commercial platforms by providing the features and ease-of-use of a proprietary software package in an extensible, open-source format. Specifically, Thunor has the following aims:

*1*) Provide a simple user interface.

*2*) Accept and store different types of cell proliferation data (viability, time course) and requisite annotation (cell type, drug, drug concentration, etc.).

*3*) Automatically fit dose–response models and calculate values of relevant derived parameters.

*4*) Provide tools for visualization of the data in multiple formats.

*5*) Allow users to add custom annotation.

*6*) Automatically perform basic statistical tests.

*7*) Be free, open-source, and extensible, allowing incorporation of modifications and features suggested by users and the research community at large.

*8*) Enable inclusion of large, pre-existing datasets.

*9*) Provide easy data access while maintaining data security to foster collaboration among research groups.

Below, we provide brief descriptions of numerous open-source and proprietary software packages for analyzing and visualizing cell proliferation data. The list is not exhaustive but, in our opinion, is representative of the myriad software platforms available to researchers in this area. Pricing and features are correct as of 29$^{th}$ January, 2021. A feature comparison is shown in Table S1.

#### Free and open-source tools

- **GRcalculator / GRmetrics**: GRcalculator (Clark et al., 2017) is an open-source, web-based graphical user interface (GUI) analogous to Thunor Web. GRmetrics is an open-source software package written in R that is analogous to Thunor Core and has been integrated with GRcalculator. These tools are based on the growth rate (GR) inhibition drug-effect metric (Hafner et al., 2016). The GR metric is closely related to the drug-induced proliferation (DIP) rate metric (Harris et al., 2016); the primary difference being that GR is defined in terms of a ratio with respect to the control rate of growth and is scaled between $-1$ (cytotoxicity) and 1 (no effect). GRcalculator allows users to upload their own data and features dose–response curve fitting, calculation of derived dose–response parameters, and various visualization tools. End-point viability (cell count relative to control) is also calculated if initial cell counts ($t = 0$) is provided for all conditions. Access to several large datasets from the Library of Integrated Network-Based Cellular Signatures (LINCS) consortium is provided, including the CTRP (but not GDSC). Further information can be found at grcalculator.org.

- **drc**: A flexible package within the R statistical programming environment for fitting dose–response curves and extracting derived parameters. It is not specific to any particular data type and can therefore be used with both viability and time-course data. A comprehensive overview of the utility of the `drc` package for the analysis of dose–response data was recently published (Ritz et al., 2015). Further information can be found at CRAN.R-project.org/package=drc.

- **PharmacoDB**: An open-source, online tool for visualizing and analyzing large pre-existing drug–response datasets (Smirnov et al., 2017), including the GDSC and CTRP. Does not have an upload capability to incorporate user-generated data. All response data are represented in terms of cell viability as a percentage of control. The resource allows users to search for instances where drugs or cell lines of interest have been profiled within one or more of various large datasets and uses the extracted dose–response parameters to identify molecular features (such as gene expression levels) associated with them. Further information can be found at pharmacodb.pmgenomics.ca.

#### Proprietary tools

- **GraphPad Prism**: Statistical software designed to perform many different types of statistical tests and generate graphs, including dose–response curves. It has a user-friendly interface that provides non-statisticians with assistance in choosing appropriate statistical tests, which has made it a very popular tool for the biomedical research community, particularly in academic settings. However, it was not designed to be a data repository with any search capabilities nor was it designed to handle the many thousands of data points present in large datasets. A single user perpetual license cost is $725 ($525 academic), but other license fee options are available when purchasing at graphpad.com.

- **IncuCyte**: IncuCyte Zoom and S3 instruments come with software to control data acquisition that is also capable of data analysis. These systems are inherently capable of acquiring time-course cell count data, including plate map information, but the software currently does not calculate proliferation rates nor can it accept data from other sources. The software has some sharing and security features—controlled access to all data on the instrument—but cannot control access of individual users to subsets of the data. The software is included with the purchase of the imaging hardware (>$110,000) but is being continually developed. Upgraded functionality may require further cost allocation. Additional information is available at essenbioscience.com/en/products/software/incucyte-base-software.

- **Genedata Screener**: Per the company's website, "Genedata Screener® analyzes, visualizes, and manages screening data from in-vitro screening assay technologies across and beyond the enterprise." The software is modular and can be tailored to the needs of the user (group). A Time Series Extension is available to accept time-course cell count data and automatically calculate proliferation rates but requires specific customization of the software. Annual license fees are in the range of $40,000–$90,000 or more. Additional information is available at genedata.com/products/screener.

- **CDD Vault**: A web-based service for storing and analyzing large drug–response datasets, primarily focused on drug development within large academic centers and corporate environments. The primary focus of the tool is to manage drug-centric data, including chemical structures and biological responses, to facilitate data exploration using graphic tools and statistical tests. Incorporation of preexisting large datasets (e.g., CTRP and GDSC) has not been described. The tool has been designed to be used collaboratively with secure and controlled access by multiple types of users, such as chemists, biologists, and data scientists. Although the breadth of capabilities available suggests that it could be extended to perform functions for which it was not initially designed, it is unclear whether CDD Vault can accept time-course cell count data and automatically calculate proliferation rates. A 30-day free trial is available but a direct consultation with Collaborative Drug Discovery, Inc. employees is required prior to obtaining the trial software or to obtain cost estimates. Additional information is available at collaborativedrug.com/cdd-vault.

### Text S3: Installation and configuration

**We recommend following the latest version of these installation instructions at docs.thunor.net**. Here, we demonstrate a typical containerized installation for evaluation or production use, but software engineers wishing to modify or extend Thunor may find the developer installation more suitable (docs.thunor.net/developer-installation).

#### Thunor Web installation

This section will show you how to install Thunor Web for testing or production use.

Thunor Web consists of several components, which are controlled together by a system called Docker Compose. Each component runs in a separate container, which makes the system modular and increases security.

A typical installation will use the following components:

- Thunor Web application server, written in Python using the Django framework.

- nginx web server, to handle HTTP requests

- Postgres database server
- Redis key-value server, used for caching time-consuming calculations and database queries.

##### Thunor Web Requirements

- Docker and Docker Compose (version 1.12 or later). Follow instructions for your platform:
  - Docker for Windows: download from docs.docker.com/docker-for-windows/install
  - Docker for Mac: download from docs.docker.com/docker-for-mac/install
  - Ubuntu 18.04 or later: sudo apt install docker-compose
  - Other platforms: see docs.docker.com/get-docker
- Python $\geq$3.6
- Git

If you use MacOS or Linux, you probably already have Python and git installed. If you use Windows, we recommend installing Anaconda (anaconda.com/download), a Python distribution which includes lots of useful software.

##### Installation steps

1. (Optional) Configure a DNS name. This is an entry like thunor.example.com which points to the machine hosting Thunor Web. If you're only testing Thunor Web on your local machine and don't need network access, this step is unnecessary.
2. Retrieve the Thunor Web quick start tool using git

   ```
   git clone https://github.com/alubbock/thunor-web-quickstart thunor-web
   cd thunor-web
   ```

3. Run the Thunor Web deploy script.

   To install a locally accessible version for testing, use:

   ```
   python thunorctl.py deploy -hostname=localhost
   ```

   To install a network accessible version at thunor.example.com, and to activate TLS encrypted connections using certbot, use:

   ```
   python thunorctl.py deploy -hostname=thunor.example.com -enable-tls
   ```

4. Create an admin account. You can use this to log in.

   ```
   python manage.py createsuperuser
   ```

That's it! Thunor Web is now ready to start.

##### Starting and stopping Thunor Web

To start Thunor Web:

```
python thunorctl.py start
```

To stop Thunor Web:

```
python thunorctl.py stop
```

These commands must be run from the thunor-web directory.

##### Enable TLS encryption

To enable TLS encryption for secure connections, you'll need a specific domain name pre-configured with a DNS A record pointing to the server IP address. Your domain registrar can help with this step.

To generate and enable TLS certificates, simply run:

```
python thunorctl.py generatecerts
```

Further information is available in the documentation at docs.thunor.net/enable-https-encryption

##### Upgrade Thunor Web

Check your current version with:

```
python thunorctl.py version
```

To upgrade to the latest version, run:

```
git pull
python thunorctl.py upgrade
```

In production, a database backup is recommended before performing an upgrade. Using the setup described above, you can just stop Thunor Web and backup the entire thunor-web directory.

Garnett, M. J., Edelman, E. J., Heidorn, S. J., Greenman, C. D., Dastur, A., Lau, K. W., Greninger, P., Thompson, I. R., Luo, X., Soares, J., Liu, Q., Iorio, F., Surdez, D., Chen, L., Milano, R. J., Bignell, G. R., Tam, A. T., Davies, H., Stevenson, J. A., Barthorpe, S., Lutz, S. R., Kogera, F., Lawrence, K., McLaren-Douglas, A., Mitropoulos, X., Mironenko, T., Thi, H., Richardson, L., Zhou, W., Jewitt, F., Zhang, T., O'Brien, P., Boisvert, J. L., Price, S., Hur, W., Yang, W., Deng, X., Butler, A., Choi, H. G., Chang, J. W., Baselga, J., Stamenkovic, I., Engelman, J. A., Sharma, S. V., Delattre, O., Saez-Rodriguez, J., Gray, N. S., Settleman, J., Futreal, P. A., Haber, D. A., Stratton, M. R., Ramaswamy, S., McDermott, U. and Benes,

C. H. (2012). Systematic identification of genomic markers of drug sensitivity in cancer cells. Nature *483*, 570–575.

Hafner, M., Niepel, M., Chung, M. and Sorger, P. K. (2016). Growth rate inhibition metrics correct for confounders in measuring sensitivity to cancer drugs. Nat. Methods *13*, 521–527.

Harris, L. A., Frick, P. L., Garbett, S. P., Hardeman, K. N., Paudel, B. B., Lopez, C. F., Quaranta, V. and Tyson, D. R. (2016). An unbiased metric of antiproliferative drug effect *in vitro*. Nat. Methods *13*, 497–500.

McKinney, W. (2010). Data structures for statistical computing in Python. In Proceedings of the 9th Python in science conference (SCIPY 2010), (van der Walt, S. and Millman, J., eds), pp. 51–56 , Austin, TX.

Merkel, D. (2014). Docker: lightweight Linux containers for consistent development and deployment. Linux J. *2014*, 2.

Ritz, C., Baty, F., Streibig, J. C. and Gerhard, D. (2015). Dose-response analysis using R. PLoS One *10*, e0146021.

Smirnov, P., Kofia, V., Maru, A., Freeman, M., Ho, C., El-Hachem, N., Adam, G.-A., Ba-alawi, W., Safikhani, Z. and Haibe-Kains, B. (2017). PharmacoDB: an integrative database for mining *in vitro* anticancer drug screening studies. Nucleic Acids Res. *46*, D994–D1002.

van der Walt, S., Colbert, S. C. and Varoquaux, G. (2011). The NumPy array: a structure for efficient numerical computation. Comput. Sci. Eng. *13*, 22–30.

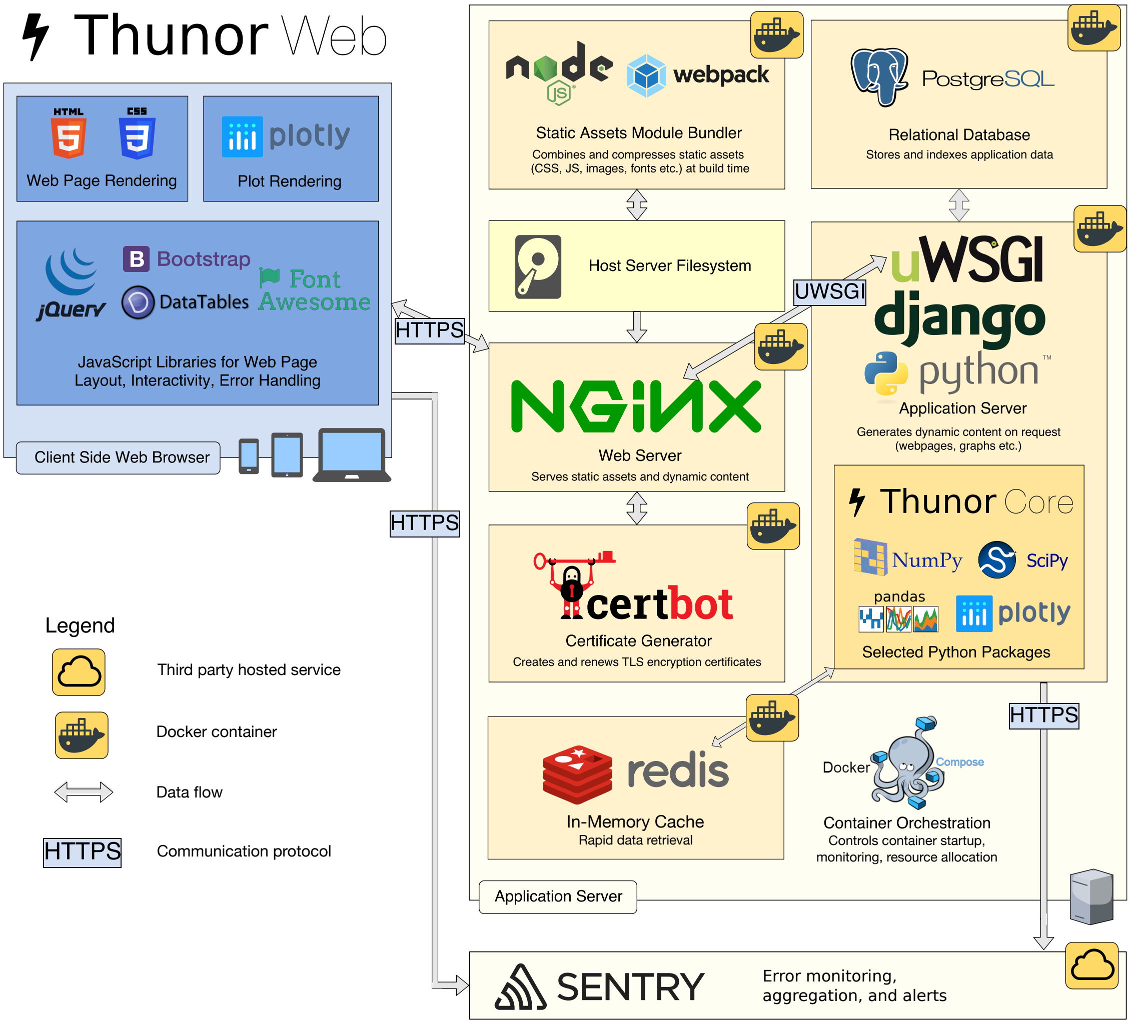


**Fig. S1. Thunor Web software infrastructure.** Users access Thunor Web through a web browser. Thunor Core produces plots using the `plotly` library, which can be delivered through Thunor Web. On the server, Thunor Web runs in multiple Docker containers: `nginx` web server, `PostgreSQL` database, and application server. The application server runs `uWSGI` and `Django`, a Python web application framework that interfaces with Thunor Core. Encryption certificates are generated using `Certbot` in a separate container, run on demand. Static files (CSS, Javascript, etc.) are compiled into bundles using `Webpack` to improve server performance. Error reporting and aggregation use the `Sentry` platform. The server system is "orchestrated" using `Docker Compose`.

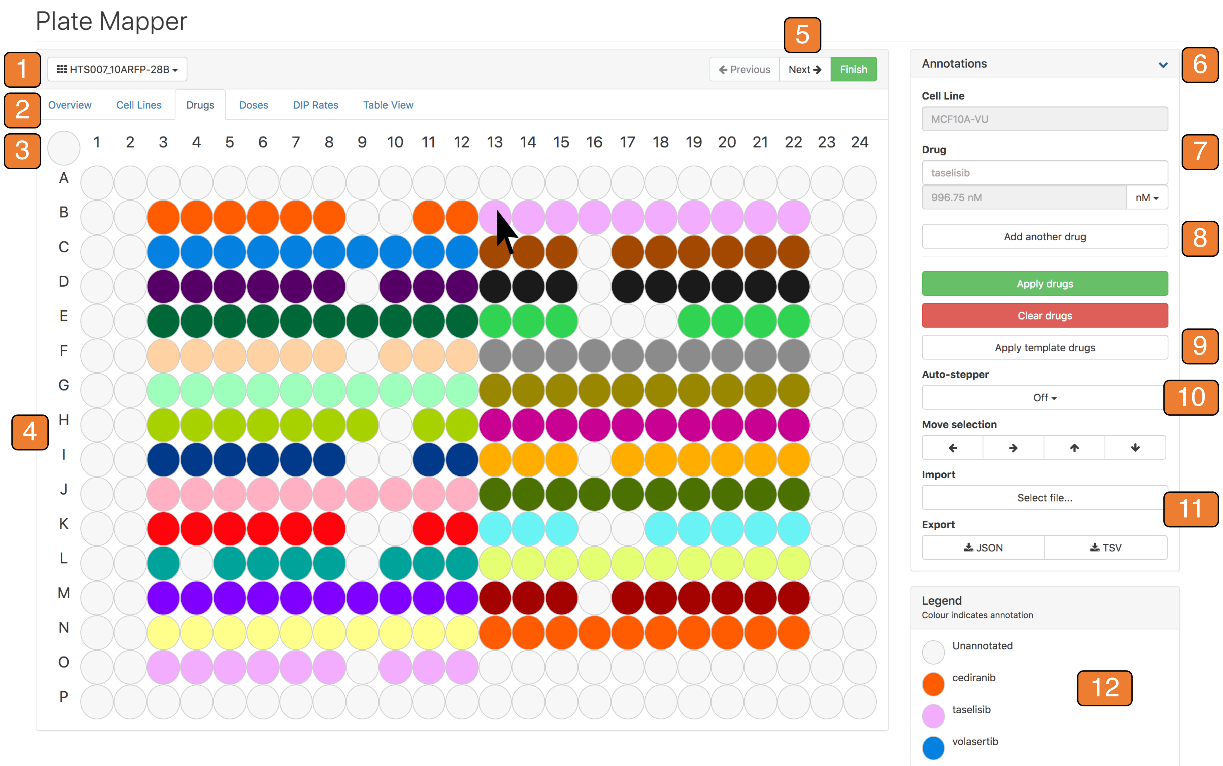


**Fig. S2. Thunor Web plate mapper.** Plates with 96, 384, and 1536 wells can be annotated and visualized using Thunor. Highlighted features of the plate mapper tool: *1.* Plate map selection menu; *2.* Change plate map colors by toggling active tab; *3.* Select all/none wells; *4.* Plate map (384 well plate shown). Color coding changes with active tab. User drags mouse/finger (touch devices) to select wells. Hover over wells to display current annotation in "Annotations" panel; *5.* Plate navigation (previous/next plate in dataset, or exit); *6.* "Annotations" panel collapse/expand toggle; *7.* Current annotation and data entry; *8.* Drug combination capability; *9.* Apply annotations from template; *10.* Auto-stepper and auto-dilution capability for rapid data entry; *11.* Plate map import and export (JSON and TSV formats); *12.* Legend for current view (cropped), changes with active tab.

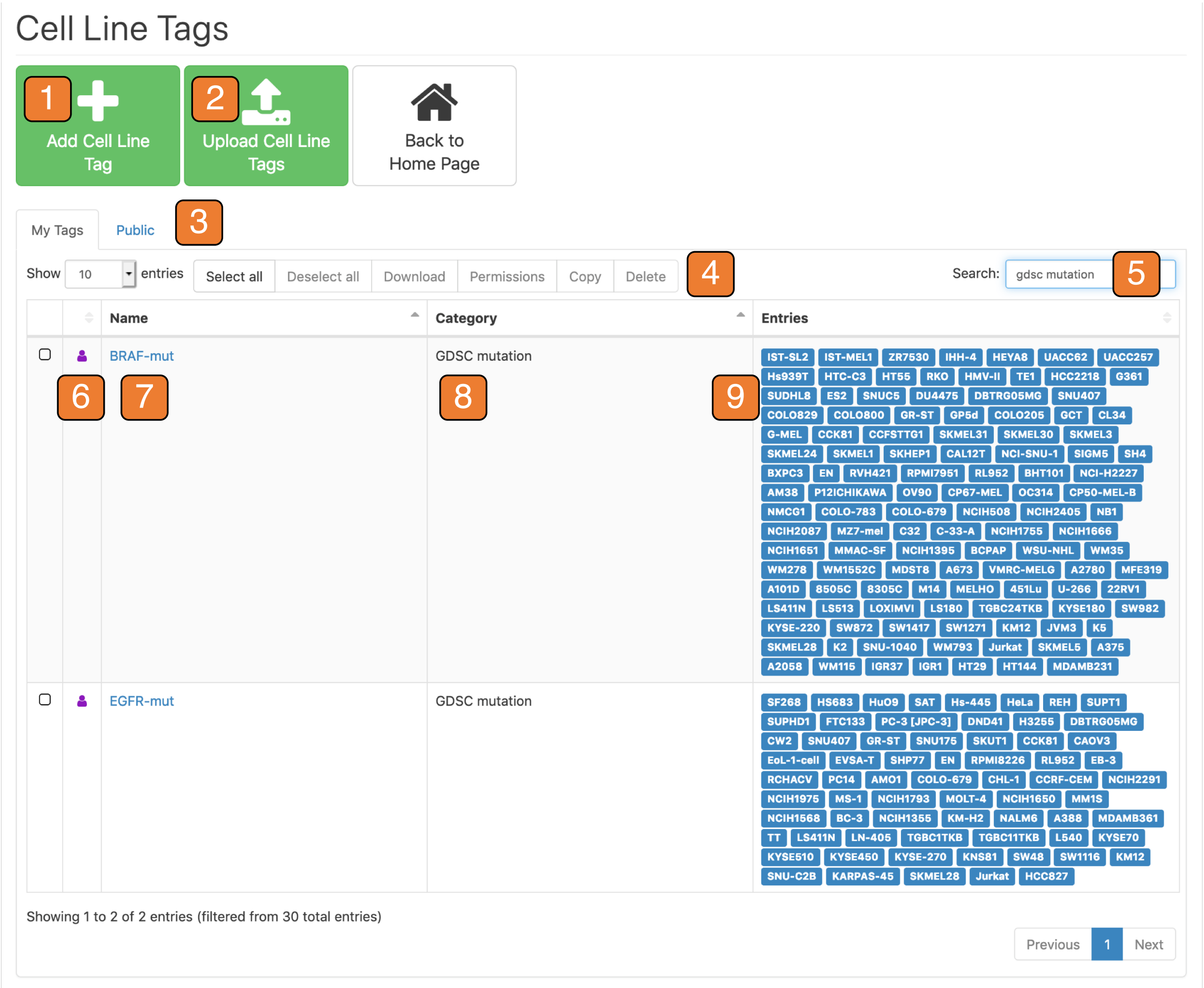


**Fig. S3. Thunor Web tag interface.** Cell line version is shown but a similar interface is available for drugs. Tags can be used to group cell lines or drugs for visualization purposes or for statistical tests. Highlighted features of the tag interface: *1.* Add tag using graphical editor; *2.* Upload tags from CSV or TSV file; *3.* View user's tags, tags by group, or public tags; *4.* Operations with selected tags: download, set group access permissions, copy (includes copy, union, and intersection), and delete; *5.* Search facility (search by tag name, category, or cell line/drug); *6.* Tag owner icon (hover cursor for details); *7.* Tags listed in paginated table—click link to edit or delete tag; *8.* Tag category (user defined); *9.* Tag entries (cell lines or drugs).

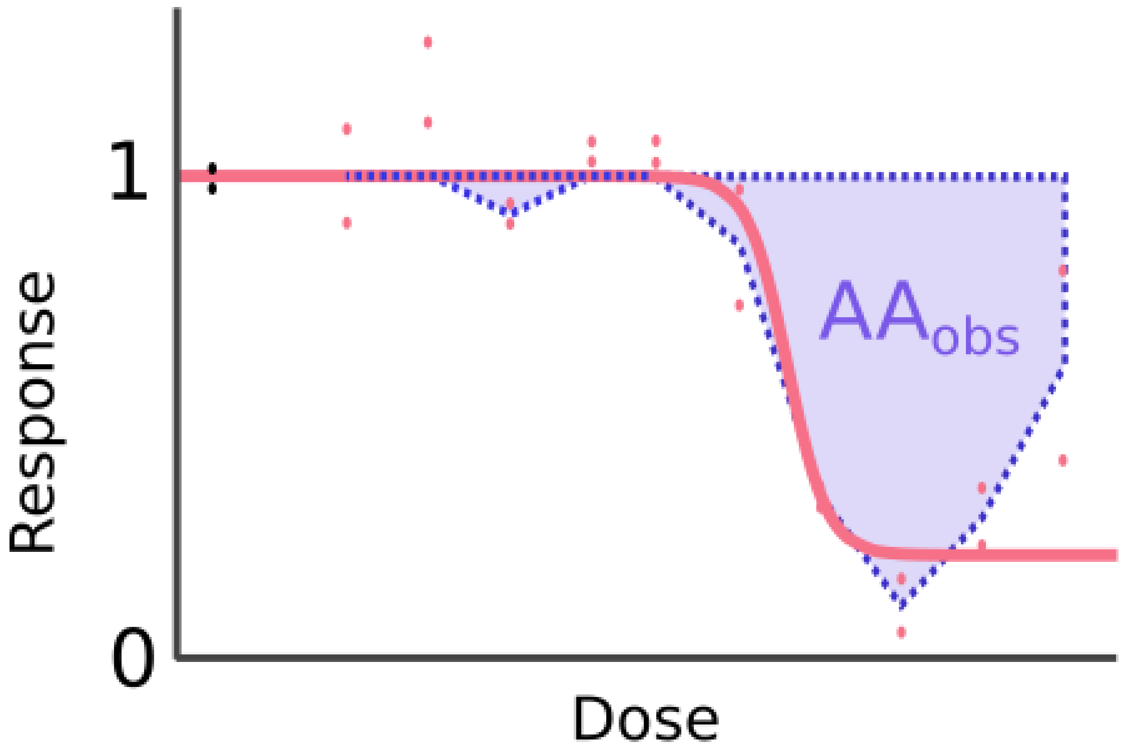


**Fig. S4. Observed activity area calculation based on observed data points (red points). Dose–response curve fit shown for comparison purposes.**

**Table S1. Feature comparisons of software for analyzing/visualizing cell proliferation data.**

| Name | Free and open source | Web interface | Upload datasets | Automatic DRC fitting | Statistical tests | Proliferation rates | Sharing and security | Tag system | Public database access | Plate annotation |
|---|---|---|---|---|---|---|---|---|---|---|
| Thunor | ✓ | ✓ | ✓ | ✓ | ✓ | ✓ | ✓ | ✓ | ✓ | ✓ |
| GRmetrics/GRcalculator[1] | ✓ | ✓ | ✓ | ✓ | | ✓ | | | ✓ | |
| drc[2] (R package) | ✓ | | ✓ | | ✓ | ✓* | | | | |
| PharmacoDB[3] | ✓ | ✓ | | ✓ | | | | | ✓ | |
| GraphPad Prism[4] | | | ✓ | | ✓ | ✓* | | | | |
| IncuCyte[5] | | | | ✓ | | | ✓ | | | ✓† |
| Genedata Screener[6] | | | ✓ | ✓ | ✓ | ✓* | ✓ | | | ✓ |
| CDD Vault[7] | | ✓ | ✓ | ✓ | ✓ | | ✓ | ✓ | | ✓ |

[1]Clark et al., *BMC Cancer* **17**, 698 (2017); grcalculator.org

[2]Ritz & Streibig, *J. Stat. Softw.* **12**, 1–22 (2005); CRAN.R-project.org/package=drc

[3]Smirnov et al., *Nucleic Acids Res.* **46**, D994–D1002 (2018); pharmacodb.pmgenomics.ca

[4]graphpad.com/prism

[5]essenbioscience.com/en/products/software/incucyte-base-software

[6]genedata.com/products/screener

[7]collaborativedrug.com/cdd-vault

*Possible but non-intrinsic feature

†Partial capability
