## Supplementary material for "Thunor: Visualization and Analysis of High-Throughput Dose-response Datasets": Data File S1: README.txt

Example Thunor Cell Line Tags
=============================

The attached file contains example tags for Thunor Web (www.thunor.net).
Tags are categorized text labels. In this case, we have provided
two example tags: "aerodigestive tract" and "skin", which are primary
tissue classifications for cell lines, based on categories in the
Genomics of Drug Sensitivty in Cancer (GDSC) project.

The attached CSV file can be uploaded in the Thunor Web tag system
by drag-and-drop. Further instructions can be found in the manual at
docs.thunor.net
