## Supplementary material for "Thunor: Visualization and Analysis of High-Throughput Dose-response Datasets": Data File S2: README.txt

HTS007 Dataset
==============

The attached HTS007 dataset contains a panel of eight breast cancer cell lines
treated with 27 drugs at multiple concentrations (4-fold dilutions). Each cell
line was modified to express fluorescent histone 2B (H2BmRFP) to enable
detection of nuclei via fluorescence microscopy[1,2]. Cells were imaged by automated
fluorescence microscopy approximately every four hours over five days in the
Vanderbilt University High Throughput Screening Core. Nuclei were quantified
by automatic image segmentation. The dataset was generated and assembled by
Darren Tyson <>.

The file is designed to be used with the Thunor Core and Thunor Web packages, which
are available at www.thunor.net. Documentation is available at the same address.

1.	Fractional proliferation: a method to deconvolve cell population dynamics from single-cell data. 9, 923–928 (2012).
2.	Trait variability of cancer cells quantified by high-content automated microscopy of single cells. 467, 23–57 (2009).
